# CRISPR/Cas9 gene editing uncovers the role of CTR1 and ROS1 in melon fruit ripening and epigenetic regulation

**DOI:** 10.1101/2022.01.30.478227

**Authors:** Andrea Giordano, Miguel Santo Domingo, Leandro Quadrana, Marta Pujol, Ana Montserrat Martín-Hernández, Jordi Garcia-Mas

## Abstract

Melon (*Cucumis melo* L.) has emerged as an alternative model to study fruit ripening due to the coexistence of climacteric and non-climacteric varieties. The previous characterization of a major QTL *ETHQV8.1* sufficient to trigger climacteric ripening in a non-climacteric background allowed the identification within the QTL interval of a negative regulator of ripening *CmCTR1*-like (MELO3C024518), and a putative DNA demethylase *CmROS1* (MELO3C024516), the orthologue of *DML2*, a DNA demethylase regulating fruit ripening in tomato. To understand the role of these genes in climacteric ripening, we generated homozygous CRISPR knockout mutants of *CmCTR1*-like and *CmROS1* in a climacteric genetic background. The climacteric behavior was altered in both loss-of-function mutants in two summer seasons with an advanced ethylene production profile compared to the climacteric wild type, suggesting a role of both genes in climacteric ripening in melon. Single cytosine methylome analyses of the *CmROS1* knockout mutant revealed DNA methylation changes in the promoter regions of key ripening genes as *ACS1*, *ETR1* and *ACO1*, and ripening associated-transcription factors as *NAC-NOR, RIN* and *CNR*, suggesting the importance of *CmROS1*-mediated DNA demethylation for triggering fruit ripening in melon.

## Introduction

During the ripening process, fleshy fruits undergo physiological and metabolic changes affecting color, flavor, firmness, and aroma. These changes are driven by phytohormones and developmental factors and occur in a highly coordinated manner with a direct impact on fruit quality and shelf-life (Giovannoni, 2001). One of the main promoters of fruit ripening is the volatile hormone ethylene. Depending on the involvement of this hormone during ripening, fruits have been traditionally divided into i) climacteric, characterized by an increase in respiration and ethylene production at the onset of ripening and ii) non climacteric, presenting low levels of both ethylene production and respiration rate across the process (McMurchie *et al*., 1972). Dissecting the regulatory network underlying the control of fruit ripening has been a major goal due to its biological significance but also for its commercial value (Giovannoni *et al*., 2017; Wang *et al*., 2020).

Important advances in the understanding of the molecular mechanisms underlying climacteric fruit ripening have been made in the model species tomato (Giovannoni, 2007) Ripening related mutants allowed the identification of several transcription factors that are upstream regulators of ethylene dependent or independent ripening. Among them *RIPENING INHIBITOR* (*RIN*), *NON-RIPENING* (*NOR*), *and COLORLESS NON-RIPENING* (*CNR*) (Vrebalov *et al*., 2002; Manning *et al*., 2006; Giovannoni, 2007).

Recent studies demonstrated that DNA methylation levels play an important role at the onset of fruit ripening in tomato(Zhong *et al*., 2013). Modulation of DNA methylation levels is governed by DNA methylases and demethylases. The enzymatic removal of methylcytosine in plants is initiated by a family of DNA glycosylases/lyases, including DEMETER (DME), Repressor of silencing 1 (ROS1), DEMETER-like2 (DML2) and DEMETER-like3 (DML3), firstly characterized in the model plant *Arabidopsis thaliana* (Zhu, 2009). In tomato, SlDML2 is induced upon the onset of ripening leading to a global DNA hypomethylation during ripening (Zhong *et al*., 2013; Liu *et al*., 2015; Lang *et al*., 2017) Knockout using CRISPR/Cas9 system and knockdown RNAi mutants in this species revealed that *SlDML2* is required for normal fruit ripening by the activation of ripening-induced genes and repression of several ripening-repressed genes (Zhong *et al*., 2013; Lang *et al*., 2017). Nonetheless, the tomato model is not universal as different transcriptional positive feedback circuits controlling ripening in climacteric species were identified (Lü *et al*., 2018).

Melon (*Cucumis melo* L.) has emerged as an alternative model to study fruit ripening since both climacteric (e.g. *cantalupensis* types as ‘Védrantais’ (VED)) and non-climacteric (e.g. *inodorus* types as ‘Piel de Sapo’ (PS)) genotypes exist. The recent characterization of a major QTL in chromosome 8 of melon, *ETHQV8.1*, which is sufficient to activate climacteric ripening in a non-climacteric background, allowed the identification of candidate genes related to fruit ripening in a genomic interval of 150 kb that contained 14 annotated genes (Pereira *et al*., 2020). Some of these genes are highly expressed in fruits and contain multiple non-synonymous polymorphisms distinguishing the climacteric VED from the non-climacteric PS genotype.

One of the candidates (*CmROS1*, MELO3C024516) encodes the homolog of the main DNA demethylase *ROS1* in Arabidopsis, which targets mainly transposable element (TE) sequences and regulates some genes involved in pathogen response and epidermal cell organization(Yamamuro *et al*., 2014; Le *et al*., 2014). The closest orthologue in tomato, *SlDML2* is crucial for the DNA demethylation of fruit ripening genes including ethylene synthesis and signaling (Lang *et al*., 2017).

The other candidate gene is *CONSTITUTIVE TRIPLE RESPONSE 1* (*CTR1*) a serine/threonine kinase (*CmCTR1-like*, MELO3C024518). This kinase interacts physically with ethylene receptors as a negative regulator of the ethylene signal transduction pathway(Kieber *et al*., 1993). In the absence of ethylene, *CTR1* is activated, preventing the downstream transduction pathway; when ethylene is present, the ethylene receptor terminates the activation of *CTR1*, leading to the ethylene responses (Binder, 2008). In tomato, the silencing of *CTR1* promoted fruit ripening, validating its role as a negative regulator of the ethylene signal transduction pathway (Fu *et al*., 2005).

In this study, we aimed to better understand the role of the two *ETHQV8.1*-containing candidate genes *CmROS1* and *CmCTR1*-like in fruit ripening by obtaining CRISPR/Cas9-induced loss-of-function mutants in a climacteric melon genotype. Furthermore, we characterized the role of *CmROS1* in DNA methylation homeostasis during fruit ripening.

## Materials and methods

### CRISPR/Cas9 vector Construction

To target *CmROS1* (MELO3C024516) two different guide RNAs (gRNA) of 20 nucleotides in length separated by 188 bp were designed using Breaking Cas tool (https://bioinfogp.cnb.csic.es/tools/breakingcas/) (Table S1). The two oligonucleotides generated for each gRNA were annealed and cloned in the sites *Bbs*I and *Bsa*I into the plasmid ptandemgRNA. The construct was verified by sequencing and then digested with *Spe*I and *Kpn*I to release the cassette that was then inserted into the same sites in the pB7-Cas9-TPC-polylinker binary vector. Cloning vectors were gently provided by Prof. Puchta (KIT, Germany).

For *CmCTR1*-like (MELO3C024518) we used the pEn-CHIMERA vector provided by Prof. Puchta (KIT, Germany) to generate the entry construct. A single gRNA of 20 nucleotides was designed using Breaking Cas tool (https://bioinfogp.cnb.csic.es/tools/breakingcas/) (Table S1). Cloning steps of the gRNA and transfer to the pDe-Cas9 binary vector were performed as previously described(Schiml and Puchta, 2016).

### Agrobacterium mediated plant transformation

*Agrobacterium tumefaciens* (strain AGL0) cells were transformed with the binary CRISPR/Cas9 construct. Plant transformation was performed by co-cultivation of the Agrobacterium culture with one-day-old cotyledons of VED as previously described (Castelblanque *et al*., 2008) including the following modifications: cotyledons were dissected as in (García-Almodóvar *et al*., 2017), regeneration media was supplemented with 6-bencylaminopurine (BA) and Indole-3-acetic acid (IAA) and *Agrobacterium* was co-cultured with the explants during a period of three days. Transgenic plants containing the *bar* gene were selected by L-Phosphinothricin (PPT) resistance and were grown in a growth room under a 12-h light/12-h dark cycle at 28 °C.

### Detection of mutations

Genomic DNA from leaves of in vitro plantlets (T0) and from young leaves of T1 and T2 plants was extracted using the CTAB method with some modifications as described in (Pereira *et al*., 2018). The transgene presence was detected by PCR using specific primers targeting Cas9. Genomic regions flanking gRNA1 and gRNA2 of *CmROS1* were amplified by PCR using specific primers. For detection of mutations in *CmCTR1*, a region targeting the gRNA was amplified with specific primers. All primers are listed in Table S2. Mutations were detected by sequencing the amplified fragments and identified by double peaks in the sequence chromatograms. Purified PCR products were cloned into p-Blunt II-TOPO vector (Life Technologies) and sequencing of colonies using M13F and M13R primers was performed to confirm the mutations.

### Generation of T2 plants and phenotyping of climacteric ripening traits

Ploidy level of T0 plants was evaluated by flow-cytometry analysis and selected T0 plants for each gene were grown under greenhouse conditions (25°C for 16 hours and 22°C for 8 hours) and self-pollinated. T1 seedlings were screened for the presence of Cas9 by PCR. After segregation, non-transgenic homozygous edited T1 plants were selected and grown under greenhouse conditions to obtain the T2 seeds for the phenotypic assay.

Edited T2 *CmROS1* (n=8) and *CmCTR1* plants (n=8) were grown randomized under greenhouse conditions (24°C for 16 hours and 22°C for 8 hours) at Caldes de Montbui (Barcelona) in 2020 and 2021. VED plants were used as a wild type control plant (n=8). Plants were weekly pruned and manually pollinated to obtain one fruit per plant. The harvest date was determined following two criteria: either abscission date, when the fruit abscised from the plant, or 5 days after the formation of the abscission layer when it was not complete.

Ripening-related traits were evaluated as described in Pereira 2020 in two consecutive summer seasons (2020 and 2021). Production of aroma (ARO), chlorophyll degradation (CD) and abscission layer formation in the pedicel of the fruit (ABS) were daily evaluated and firmness was measured at harvest time. The visual inspection of melon fruits, attached to the plant, was performed daily, from approximately 20 days after pollination (DAP) until harvest. In addition, individual pictures of the fruits were obtained weekly. ARO, ABS and CD were recorded as 0 = absence and 1 = presence. The aroma production was evaluated every day by smelling the fruits. The firmness of fruit flesh was measured at harvest using a penetrometer (Fruit Test™, Wagner Instruments), in at least three regions of the fruit (distal, proximal and median), and the mean value was registered.

### Ethylene production

Ethylene production *in planta* was measured in the 2020 summer season using non-invasive gas chromatography – mass spectrometry (GC-MS) method, as described in (Pereira *et al*., 2017). The ethylene peak was monitored before ripening from 20 DAP until harvest. The atmosphere of the chamber containing the fruit was measured every day.

The ethylene peak was characterized by four traits, measured as described in Pereira 2020: maximum production of ethylene in the peak (ETH), earliness of ethylene production (DAPE), earliness of the ethylene peak (DAPP), and width of ethylene peak (WEP).

### Epigenomics

DNA was extracted from fruit flesh of ROS1-CRISPR-2 and the wild-type VED at different ripening stages (15, 25 and 30 DAP and harvest point) following the CTAB protocol(Doyle and JJ, 1990) adding a purification step using Phenol:Chloroform:Isoamyl alcohol (25:24:1). For each time point, three biological replicates were analysed. Bisulfite conversion, BS-seq libraries and sequencing (paired-end 100 nt reads) were performed by BGI Tech Solutions (Hong Kong). Mapping was performed on melon genome v3.6.1 (Ruggieri *et al*., 2018) using Bismark v0.14.2 (Krueger and Andrews, 2011) and the parameters: --bowtie2, -N 1, -p 3 (alignment); --ignore 5 --ignore_r2 5 -- ignore_3prime_r2 1 (methylation extractor). Only unique mapping reads were retained. The methylKit package v0.9.4 (Akalin *et al*., 2012) was used to calculate differential methylation in 100 bp non-overlapping windows (DMRs). Significance of calculated differences was determined using Fisher’s exact test and Benjamin-Hochberg (BH) adjustment of p-values (FDR<0.05) and methylation difference cutoffs of 40% for CG, 20% for CHG and 20% for CHH. Differentially methylated windows within 100bp of each other were merged to form larger DMRs. 100bp windows with at least six cytosines covered by a minimum of six (CG and CHG) and ten (CHH) reads per comparison were considered.

### Statistical analyses

All the statistical analyses and graphical representations were obtained using the software R v3.2.3 (R Development Core Team, 2011) with the RStudio v1.0.143 interface (*RStudio: Integrated development environment for R*, 2012).

## Results

### Generation of CRISPR/Cas9 knockout mutants in candidate genes for *ETHQV8.1* and inheritance of the editions

To investigate the role of Cm*ROS1* (MELO3C024516) and *CmCTR1*-like (MELO3C024518) genes in the fruit ripening process in melon, we knocked them out using the CRISPR/Cas9 gene editing system in a climacteric genetic background (VED).

A strategy with two target sites in exon 2 was used for *CmROS1* (Fig. 1). We obtained 15% transformation efficiency, recovering in total 59 transgenic rooted plants. From the transgenic plants, almost half of them (46%) were edited. Multiple independent transgenic plants were genotyped by sequencing the genomic DNA spanning both target sites. Most of the editions (75%) occurred in target 1 (gRNA1) whereas only a few editions (25%) were obtained for target 2 (gRNA2). Several different insertions and deletions were obtained in T0 plants with biallelic or heterozygous mutations (Fig. S1), with several plants carrying the same mutation (+1 bp). A diploid biallelic line with an insertion of 1 bp and a deletion of 23 bp that were predicted to generate truncated proteins was selected for further work (Fig. 1A).

**Figure 1:**
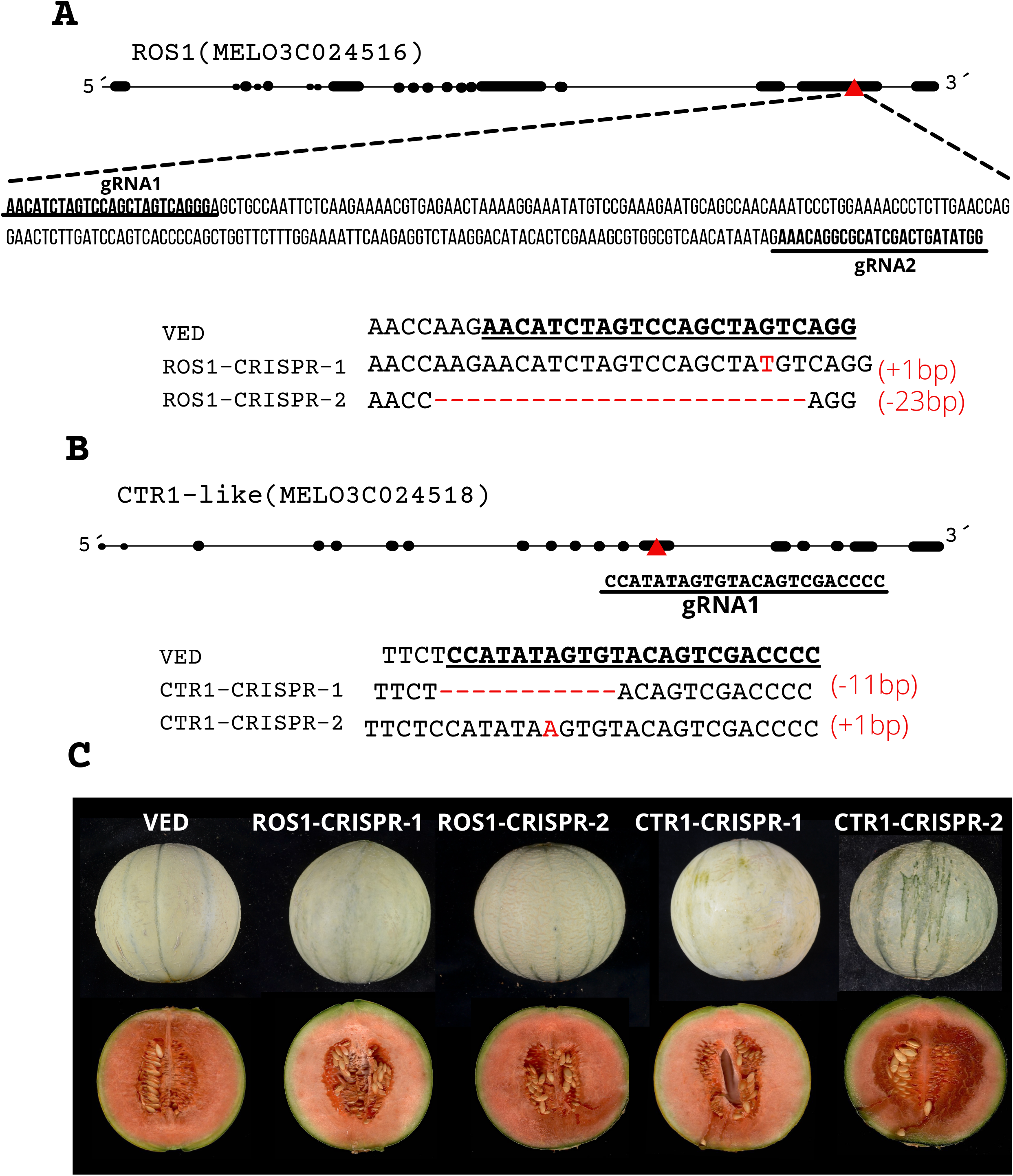
Schematic representation of the target sites for CRISPR/Cas9 and selected CRISPR edited lines. **(A)** Position of the gRNA target sites (red triangle) and selected mutations in the CRISPR lines for *CmROS1* **(B)** Position of the gRNA target sites (red triangle) and selected mutations in the CRISPR lines for *CmCTR1*-like (**C)** Fruit phenotype at harvest time of the wild type VED and CRISPR edited lines.

The selected biallelic T0 line was self-pollinated to obtain non-transgenic (Cas9 free) plants carrying homozygous editions. After segregation, T1 lines homozygous for the 1 bp insertion (ROS1-CRISPR-1) or the 23 bp deletion (ROS1-CRISPR-2) were selected for further study (Fig. 1A and C).

A different CRISPR Cas9 strategy was used to target the *CmCTR1*-like gene. A single target site was selected in exon 6 of *CmCTR1*-like (Fig. 1B). For this target gene we obtained 12% of transformation efficiency. Transgenic T0 plants were screened for mutations in the target site and 40% were edited showing mainly large or small deletions (Fig. S1). From the edited T0 plants, a biallelic line carrying a 11 bp deletion and a 1 bp insertion was selected and self-pollinated to segregate out the Cas9 transgene. The genetic editions were stably transmitted to T1 plants. After segregation, a homozygous edited line carrying the 11 bp deletion (CTR1-CRISPR-1), which is predicted to generate a premature termination codon, and the homozygous line with 1 bp insertion (CTR1-CRISPR-2), generating a frame shift, were grown under greenhouse conditions for the characterization of fruit ripening related traits (Fig. 1B and C).

### *CmROS1* and *CmCTR1* edited plants show altered ripening phenotypes

ROS1-CRISPR-1/2 and CTR1-CRISPR-1/2 were evaluated and characterized for ripening related traits in two consecutive summer seasons (2020 and 2021). However, the line CTR1-CRISPR-2 was only characterized in 2021 due to a powdery mildew infection of some replicates in 2020 that prevented its evaluation. Overall, the fruit appearance (shape, weight and colour) of the CRISPR edited lines did not show major differences with the wild-type VED at harvest time and no significant changes were detected in the flesh firmness (Fig. 1C, Table S3). To better characterize the ripening process, we measured ethylene production *in planta* in 2020 with a non-invasive methodology allowing observing the phenotype of the downstream effects of this hormone.

The phenotypic characterization revealed a significant earliness of the climacteric symptoms for all the edited lines showing the same ripening behaviour pattern in both years (Fig. 2, Table S3). In 2020, the earliest climacteric symptom was sweet aroma production (EARO), which in the CRISPR edited lines for both genes appeared around two days before VED. The initiation of the rind color change, which is attributed to chlorophyll degradation (ECD), was appreciated almost simultaneously with the detection of the abscission layer formation (EALF) and both ripening-related traits arose in both *CmROS1* edited lines two days before VED. The CTR1-CRISPR-1 edited line exhibited the earliness of the ripening related traits all at the same time, which differed significantly from VED, arising around three days before than VED for ECD and EALF and two days for EARO.

**Figure 2:**
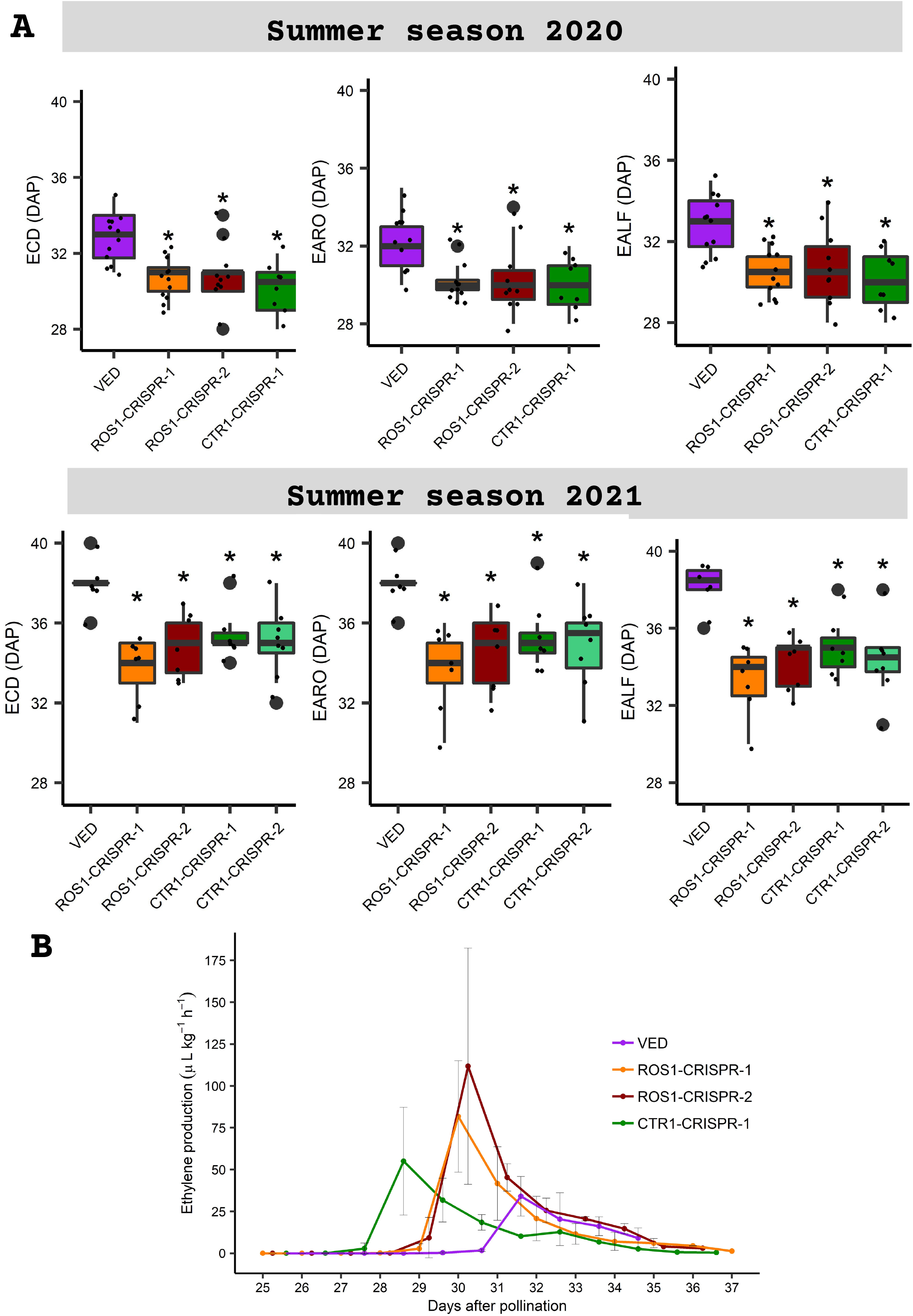
Evaluation of climacteric ripening associated traits in CRISPR edited lines and VED (in two consecutive years) and ethylene emission rates. **(A)** Earliness of chlorophyll degradation (ECD), Earliness of production of aroma (EARO) and Earliness of abscission layer formation (EALF) in 2020 **(B)** ECD, EARO and EALF in 2021 **(C)** Ethylene production in attached fruits from 25 days after pollination (DAP) until harvest in 2020.

During the second summer season, we evaluated all the CRISPR edited lines. In general, the environmental conditions delayed ripening of both VED and mutant plants (around 4-5 days later in 2021). Despite this environmental effect, all CRISPR edited lines displayed significant advances of about three days in the ripening-related traits ECD, EARO and EALF when compared to VED (Fig. 2). Moreover, during this year, the line CTR1-CRISPR-2 was evaluated, and the dataset showed the same behaviour for both CTR1 edited lines. ROS1 edited lines also showed the same pattern between them.

We also monitored fruit ethylene emission daily in 2020 without altering the ripening process (Fig. 2C). The CRISPR edited lines showed a different ethylene production pattern compared to wild-type VED, with both *CmROS1* edited lines showing the same profile. In *CmROS1* mutant lines, ethylene production started two days before the wild-type VED and with an increment of 2.7 to 3-fold of ethylene production (Fig. 2C and Table S3).

For *CmCTR1* edited lines, ethylene measurements for CTR1-CRISPR-2 were not available due to the infection with powdery mildew of some of the replicates of this line at around 20 DAP. The CTR1-CRISPR1 line showed a significant difference in the earliness of ethylene production (DAPE) and earliness of ethylene peak (DAPP). In this line, ethylene was detected around three days in advance of wild-type VED. Similarly, the peak of ethylene production was also advanced three days in CTR1-CRISPR1 compared to wild-type VED, however this advancement was not accompanied by a significant difference in the maximum quantity of ethylene produced (Fig. 2C and Table S3). Overall, these results demonstrate that both candidate genes are involved in melon fruit ripening.

### Characterization of the ROS1-CRISPR and VED methylome at different fruit ripening stages

To better understand at the molecular level the role of *CmROS1* in DNA demethylation and fruit ripening in melon, we generated single-cytosine resolution methylomes by whole genome bisulfite sequencing from fruits of ROS1-CRISPR-2 and the wild-type VED plants at 15, 25 and 30 DAP as well as at harvest (H) point (Fig. 3A).

**Figure 3:**
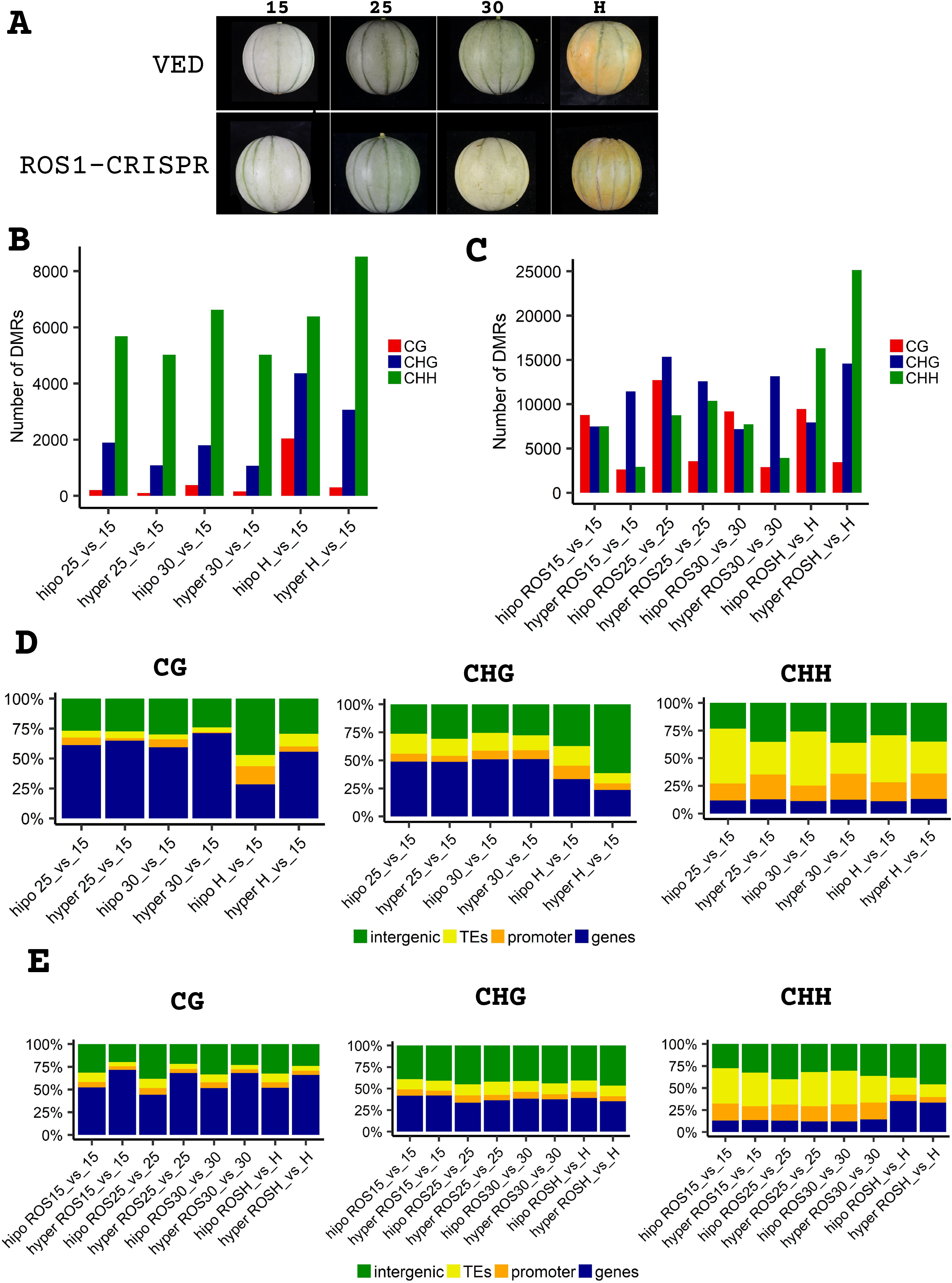
General methylation and DMR regions at different ripening stages of VED and CRISPR-ROS1 line. **(A)** Fruit ripening stages **(B)** number of DMRs along ripening in VED **(C)** number of DMRs in VED vs CRISPR-ROS1 at the same time point of ripening **(D)** DMRs annotation in VED along ripening **(E)** DMRs annotation in VED vs CRISPR-ROS1 at the same time point of ripening

When comparing the global methylation level along ripening in VED, we found that methylation at CG and CHG contexts declines along fruit ripening, showing around 2,000 and 4,000 hypomethylated regions (DMR) respectively in the CG and CHG context at harvest time compared to the first stage of ripening (i.e. H vs 15 DAP) (Fig. 3B). Interestingly, these changes were more often associated with promoter and intergenic regions (Fig. 3D).

In order to evaluate the role of ROS1 in the observed DNA methylation dynamics, we compared the methylation level in the three contexts of ROS1-CRISPR-2 and the wild-type VED plants at the same ripening stage (Fig. 3C). In this way, we identified numerous changes in DNA methylation levels for the three sequence contexts. In total (CG, CHG and CHH context together), we found 16,968 hypermethylated DMRs at 15 DAP, 26,497 at 25 DAP, 19,928 at 30 DAP and 43,156 at H time relative to VED. Overall, CRISPR-ROS1 line is associated with hypomethylation of CG and hypermethylation of CHG DMRs (Fig. 3C).

To further investigate the targets of ROS1 we focused on the hypermethylated DMRs in the CRISPR-ROS1 line (Table S4). Moreover, in CHH context at H time there are changes in the number of DMRs annotation between VED and the edited line. Among the CHH hypermethylated regions in CRISPR-ROS1 compared to VED at H time, 14% are associated with TEs, 46% with intergenic, 6% in promoter regions (defined as 1 kb upstream transcriptional start sites), and 33% in genic regions (Fig. 3D).

Notably, Gene ontology (GO) enrichment analysis of genes associated with hypermethylated DMRs at H time in the CRISPR-ROS line compared to VED and hypomethylated along ripening in VED context revealed an overrepresentation of genes related to response to stress in CRISPR-ROS1 compared to VED (Table S5).

### *CmROS1* targets promoter regions of key genes involved in ripening

We have further analysed the methylation level of key genes known to participate in the ripening process in the three contexts. Changes were found at different stages of ripening in the promoter region of genes involved in the ethylene biosynthesis or signaling pathway: *ACS1* (MELO3C016340.2), *ETR1* (MELO3C003906.2) and *ACO1* (MELO3C014437) as well as in ripening associated-transcription factors: *NAC-NOR* (MELO3C016540), *RIN* (MELO3C026300.2) and *CNR* (MELO3C002618.2) (Fig. 4).

**Figure 4:**
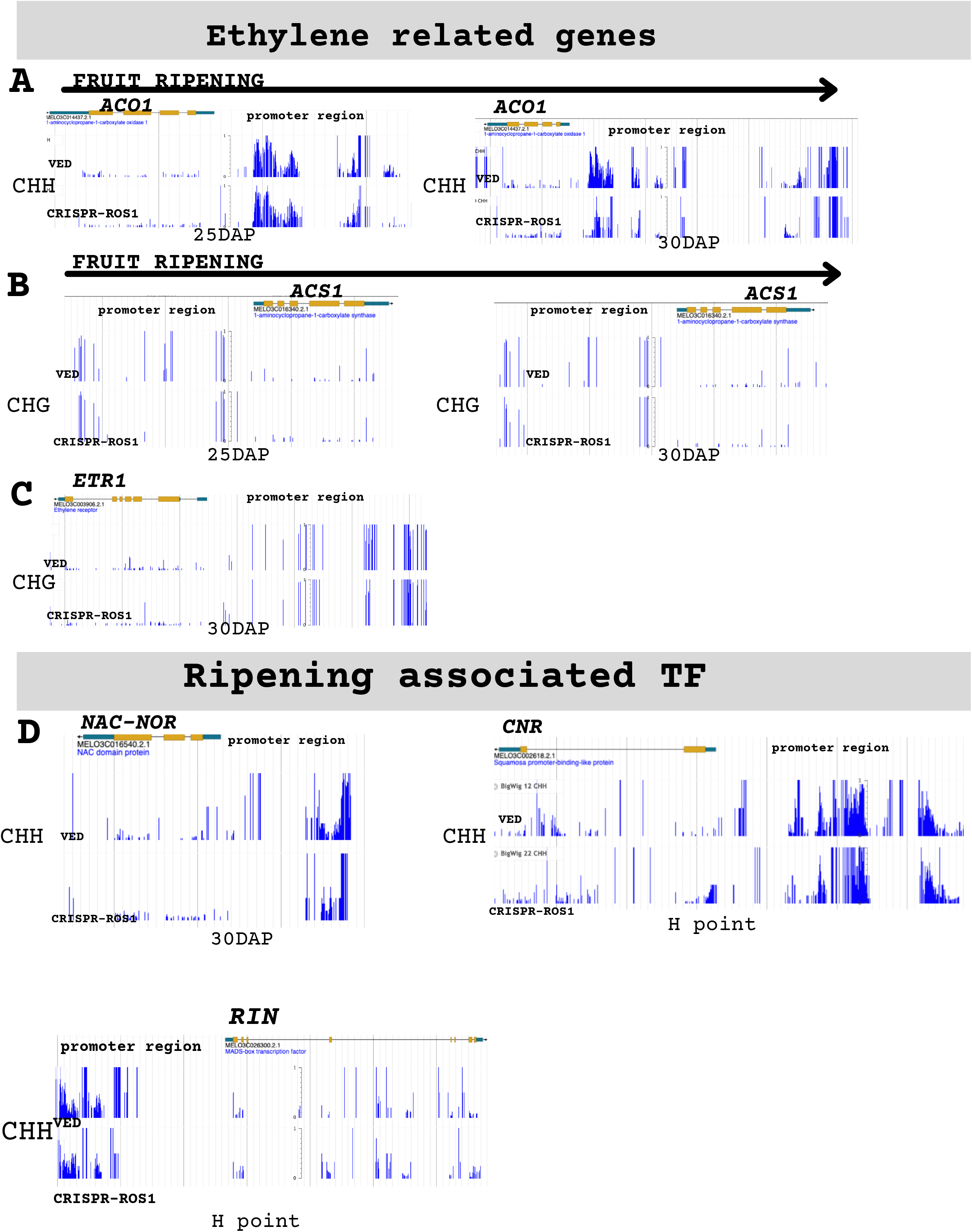
DNA methylation levels of ethylene related genes and ripening associated transcription factors for VED and CRISPR-ROS1 at different fruit ripening development stages in the three contexts **(A)** *ACO1* in CHH context at 25 (left) and 30 DAP (right) **(B)** *ACS1* in CHG context at 25 (left) and 30 DAP (right) **(C)** *ETR1* in CHG context at 30 DAP **(D)** ripening associated transcription factors in CHH context: *NAC-NOR* at 30 DAP, *RIN* at H point and *CNN* at H point.

Notably, the promoter region of *ACS1* appeared hypomethylated on the three sequence contexts in the CRISPR-ROS1 line compared to VED in all the time points studied along ripening. Furthermore, hypomethylation of the ACO1 promoter (CG and CHH context) was observed at 25 and 30 DAP and the *ETR1* promoter region (CHG and CHH context) at 30 DAP. In contrast, CHG hypermethylation of *NAC-NOR* was found from the earliest stage until 30 DAP in the mutant and hypomethylated at 30 DAP in the CHH context. For the other two transcription factors, we observed CHH hypomethylation of *RIN* and *CNR* promoter regions at H time. In combination, these results provide strong evidence that Cm*ROS1* modulates DNA methylation levels of ripening genes in melon, which may have major consequences for fruit ripening.

## Discussion

Advances in genome editing have been obtained applying the CRISPR/Cas technology in several plant species. However, among the Cucubitaceae family studies were only reported in watermelon for herbicide resistance (Tian *et al*., 2016, 2018) and cucumber for virus resistance (Chandrasekaran *et al*., 2016). More recently, edited plantlets with a disruption of a visual reporter gene (*CmPDS*), which could not be carried to the next generation, were generated in melon using CRISPR/Cas9 (Hooghvorst *et al*., 2019). To our knowledge, hereby we report for the first time the generation of melon knockout mutants for an agronomic important trait such as fruit ripening and the inheritance of the introduced mutations to the following generations using CRISPR/Cas9.

Melon is considered a recalcitrant species for genetic transformation. In this study, we obtained on average 15% transgenic plants and from these, 40% and 46% were successfully edited plants for our target genes *CmROS1* and *CmCTR1* using two or one gRNA strategy, respectively. The edited plants carried several types of editions nearby the protospacer adjacent motif (PAM) sequence of the target gRNA. As reported for other species (Feng *et al*., 2014), biallelic edited plants were obtained (70% of the edited plants), suggesting early editions during developmental stages.

In accordance with the mutations induced by Non-homologous end Joining pathway, the sequence analysis of the edited lines revealed that the most frequent editions were insertions and deletions with more than one independent event exhibiting the same edition. All the gRNA used here successfully induced mutations in the target genes. However, editions in Cm*ROS1* were mainly obtained in gRNA1 suggesting a higher edition efficiency for this gRNA. In addition, in contrast to the observations reported by Hooghvorst et. al., base pair substitutions were not obtained for any of the genes targeted in this study. Improving fruit quality and shelf life has been one of the main challenges for agriculture. During the last decades, advances in understanding the ripening process were approached by conventional breeding and genetic engineering tools. For instance, CRISPR knockout mutants in tomato have proved the importance of master ripening regulator genes(Ito *et al*., 2015).

More recent studies showed that epigenetic regulation plays a key role in fruit ripening (Lü *et al*., 2018). The balance of global DNA methylation/demethylation is altered during fruit ripening and these alterations are governed by DNA demethylases. In tomato, more than 200 promoters of ripening-related genes, including master regulators, ethylene related genes, fruit softening, and carotenoids synthesis genes, are regulated by DNA demethylation at the onset of ripening (Zhong *et al*., 2013).

In Arabidopsis, the protein repressor of silencing 1 (*ROS1*), which belongs to the subfamily of bifunctional 5-methylcytosine DNA glycosylases/lyases, has been characterized as the main sporophytic DNA demethylase(Gong *et al*., 2002). In tomato, there are four genes (*SlDML1, SlDML2, SlDML3 and SlDML4*) encoding putative DNA demethylases, being *SlDML2* the closest ortholog to Arabidopsis *ROS1* gene. Furthermore, *SlDML2* expression is highly correlated with fruit ripening (Zhong *et al*., 2013; Liu *et al*., 2015). In melon, we have identified four putative *ROS1* homologues (MELO3C024516, MELO3C021451, MELO3C002241 and MELO3C009432) (Fig S2). The gene MELO3C024516 locates in the previously identified ripening QTL interval *ETHQV8.1* and therefore was edited in this study. The CRISPR-ROS1/2 lines, carrying loss-of-function homozygous alleles of MELO3C024516, showed an advance in climacteric ripening compared to the wild type, suggesting a role of this gene in the complex regulation of climacteric ripening in melon. Interestingly, RNA-seq expression analysis of several fruit ripening stages in wild type climacteric VED shows that the four putative ROS1 genes have a similar expression profile along ripening (Fig. S3), suggesting that more than one DNA demethylase may be involved in this process. Unlike in tomato, in which the CRISPR SlDML2 mutant showed an inhibitory effect on fruit ripening (Lang *et al*., 2017), the *CmROS1* knockout melon fruit ripens ahead of the wild type VED. Our methylome analysis of the climacteric variety VED showed an overall demethylation in CG and CHG context along fruit ripening, similar to what has been reported in tomato (Liu *et al*., 2015; Lang *et al*., 2017), orange (Huang *et al*., 2019) and strawberry(Cheng et al., 2018). Furthermore, hypomethylation levels in the promoter regions of key ripening genes (e.g. *ACS1, ETR1, ACO1*) are in agreement with the phenotype displayed by the *CmROS1* CRISPR lines. Also, genes related to biotic stress response were also found to be hypomethylated in ROS1 vs VED at harvest, suggesting a possible role of this DNA demethylase in stress-response genes, as reported for *ROS1, DML2, DML3* in response to biotic stress in Arabidopsis (Le *et al*., 2014; Halter *et al*., 2021).

Both mutant lines of CTR1-CRISPR promote fruit ripening in melon in agreement to the phenotype described when silencing *LeCTR1* in tomato fruits (Fu *et al*., 2005) and the previously described role of *CTR1* as a negative regulator of ethylene signaling in other species (Binder, 2008). This second candidate gene of the QTL *ETHQV8.1* plays also an important role in the ripening process as a negative regulator affecting the initiation of the ripening process but without affecting other important traits such as firmness.

To our knowledge, this is the first time that the CRISPR technology has been implemented on genes involved in agronomically important traits in melon. The implementation of this technology in this species and the inheritance of the editions to the following generations is of high interest and a valuable resource not only for researchers but also for breeders. We have functionally validated two genes involved in the complex regulation of fruit ripening and studied in depth the role of the DNA demethylase ROS1 in fruit ripening. However, as mutants for both candidate genes *CmROS1* and *CmCTR1-like* showed an altered ripening phenotype, further studies are needed to identify which of them is the candidate for *ETHQV8.1*.

## Acknowledgements

We acknowledge F. Garcia for technical support. This work was supported by the Spanish Ministry of Economy and Competitiveness Grant RTI2018-097665-B-C2, the Spanish Ministry of Science and Innovation-State Research Agency (AEI) Severo Ochoa Programme for Centres of Excellence in R&D CEX2019-000902 and the CERCA Programme/Generalitat de Catalunya to J.G.-M. AG was supported by the European Union’s Horizon 2020 research and innovation programme under Marie Skłodowska-Curie grant agreement No 793090. MSD was supported by a FPI grant from the Spanish Ministry of Economy and Competitiveness. Work in the Quadrana group is supported by the European Research Council (ERC) under the European Union’s Horizon 2020 research and innovation program (grant agreement No. 948674).

## References

Akalin A, Kormaksson M, Li S, Garrett-Bakelman FE, Figueroa ME, Melnick A, Mason CE. 2012. MethylKit: a comprehensive R package for the analysis of genomewide DNA methylation profiles. Genome Biology 13, 1–9.

Binder BM. 2008. The ethylene receptors: Complex perception for a simple gas. Plant Science 175, 8–17.

Castelblanque L, Marfa V, Claveria E, I M, L P-G, Dolcet-sanjuan R. 2008. Improving the genetic transformation efficiency of Cucumis melo subsp. melo “Piel de Sapo” via Agrobacterium. Transformation, 627–632.

Chandrasekaran J, Brumin M, Wolf D, Leibman D, Klap C, Pearlsman M, Sherman A, Arazi T, Gal-On A. 2016. Development of broad virus resistance in non-transgenic cucumber using CRISPR/Cas9 technology. Molecular Plant Pathology, 1–14.

Cheng J, Niu Q, Zhang B, Chen K, Yang R, Zhu JK, Zhang Y, Lang Z. 2018. Downregulation of RdDM during strawberry fruit ripening. Genome Biology 19.

Doyle, JJ. 1990. Isolation of plant DNA from fresh tissue. Focus 12, 13–15.

Feng Z, Mao Y, Xu N, et al. 2014. Multigeneration analysis reveals the inheritance, specificity, and patterns of CRISPR/Cas-induced gene modifications in Arabidopsis. Proceedings of the National Academy of Sciences 111, 4632–4637.

Fu DQ, Zhu BZ, Zhu HL, Jiang WB, Luo YB. 2005. Virus-induced gene silencing in tomato fruit. Plant Journal 43, 299–308.

García-Almodóvar RC, Gosalvez B, Aranda MA, Burgos L. 2017. Production of transgenic diploid Cucumis melo plants. Plant Cell, Tissue and Organ Culture (PCTOC) 2017 130:2 130, 323–333.

Giovannoni J. 2001. MOLECULAR BIOLOGY OF FRUIT MATURATION AND RIPENING. Annual review of plant physiology and plant molecular biology 52, 725–749.

Giovannoni JJ. 2007. Fruit ripening mutants yield insights into ripening control. Current Opinion in Plant Biology 10, 283–289.

Giovannoni J, Nguyen C, Ampofo B, Zhong S, Fei Z. 2017. The Epigenome and Transcriptional Dynamics of Fruit Ripening. https://doi.org/10.1146/annurev-arplant-042916-040906 68, 61–84.

Gong Z, Morales-Ruiz T, Ariza RR, Roldán-Arjona T, David L, Zhu J-K. 2002. ROS1, a Repressor of Transcriptional Gene Silencing in Arabidopsis, Encodes a DNA Glycosylase/Lyase. Cell 111, 803–814.

Halter T, Wang J, Amesefe D, Lastrucci E, Charvin M, Rastogi MS, Navarro L. 2021. The arabidopsis active demethylase ros1 cis-regulates defense genes by erasing dna methylation at promoter-regulatory regions. eLife 10, 1–62.

Hooghvorst I, López-Cristoffanini C, Nogués S. 2019. Efficient knockout of phytoene desaturase gene using CRISPR/Cas9 in melon. Scientific Reports 2019 9:1 9, 1–7.

Huang H, Liu R, Niu Q, Tang K, Zhang B, Zhang H, Chen K, Zhu JK, Lang Z. 2019. Global increase in DNA methylation during orange fruit development and ripening. Proceedings of the National Academy of Sciences of the United States of America 116, 1430–1436.

Ito Y, Nishizawa-Yokoi A, Endo M, Mikami M, Toki S. 2015. CRISPR/Cas9-mediated mutagenesis of the RIN locus that regulates tomato fruit ripening. Biochemical and Biophysical Research Communications 467, 76–82.

Kieber JJ, Rothenberg M, Roman G, Feldmann KA, Ecker JR. 1993. CTR1, a negative regulator of the ethylene response pathway in arabidopsis, encodes a member of the Raf family of protein kinases. Cell 72, 427–441.

Krueger F, Andrews SR. 2011. Bismark: a flexible aligner and methylation caller for Bisulfite-Seq applications. Bioinformatics 27, 1571–1572.

Lang Z, Wang Y, Tang K, Tang D, Datsenka T, Cheng J, Zhang Y, Handa AK, Zhu JK. 2017. Critical roles of DNA demethylation in the activation of ripening-induced genes and inhibition of ripening-repressed genes in tomato fruit. Proceedings of the National Academy of Sciences of the United States of America 114, E4511–E4519.

Le T-N, Schumann U, Smith NA, et al. 2014. DNA demethylases target promoter transposable elements to positively regulate stress responsive genes in Arabidopsis. Genome Biology 2014 15:9 15, 1–18.

Liu R, How-Kit A, Stammitti L, et al. 2015. A DEMETER-like DNA demethylase governs tomato fruit ripening. Proceedings of the National Academy of Sciences of the United States of America 112, 10804–10809.

Lü P, Yu S, Zhu N, et al. 2018. Genome encode analyses reveal the basis of convergent evolution of fleshy fruit ripening. Nature Plants 4, 784–791.

Manning K, Tör M, Poole M, Hong Y, Thompson AJ, King GJ, Giovannoni JJ, Seymour GB. 2006. A naturally occurring epigenetic mutation in a gene encoding an SBP-box transcription factor inhibits tomato fruit ripening. Nature Genetics 2006 38:8 38, 948–952.

McMurchie EJ, McGlasson WB, Eaks IL. 1972. Treatment of fruit with propylene gives information about the biogenesis of ethylene. Nature 237, 235–6.

Pereira L, Pujol M, Garcia-mas J, Phillips MA. 2017. Non-invasive quantification of ethylene in attached fruit headspace at 1 p. p. b. by gas chromatography – mass spectrometry., 172–183.

Pereira L, Ruggieri V, Pérez S, Alexiou KG, Fernández M, Jahrmann T, Pujol M, Garcia-Mas J. 2018. QTL mapping of melon fruit quality traits using a high-density GBS-based genetic map. BMC Plant Biology 18, 1–17.

Pereira L, Santo Domingo M, Ruggieri V, et al. 2020. Genetic dissection of climacteric fruit ripening in a melon population segregating for ripening behavior. Horticulture Research 7.

R Development Core Team. 2011. A Language and Environment for Statistical Computing., 409.

RStudio: Integrated development environment for R. 2012. Boston, MA, USA: RStudio Inc.

Ruggieri V, Alexiou KG, Morata J, et al. 2018. An improved assembly and annotation of the melon (Cucumis melo L.) reference genome. Scientific reports 8.

Schiml S, Puchta H. 2016. Revolutionizing plant biology□: multiple ways of genome engineering by CRISPR / Cas. Plant Methods, 1–9.

Tian S, Jiang L, Cui X, et al. 2018. Engineering herbicide-resistant watermelon variety through CRISPR/Cas9-mediated base-editing. Plant Cell Reports 2018 37:9 37, 1353–1356.

Tian S, Jiang L, Gao Q, et al. 2016. Efficient CRISPR/Cas9-based gene knockout in watermelon. Plant Cell Reports 2016 36:3 36, 399–406.

Vrebalov J, Ruezinsky D, Padmanabhan V, White R, Medrano D, Drake R, Schuch W, Giovannoni J. 2002. A MADS-box gene necessary for fruit ripening at the tomato ripening-inhibitor (rin) locus. Science (New York, N.Y.) 296, 343–346.

Wang R, Angenent GC, Seymour G, de Maagd RA. 2020. Revisiting the Role of Master Regulators in Tomato Ripening. Trends in Plant Science 25, 291–301.

Yamamuro C, Miki D, Zheng Z, Ma J, Wang J, Yang Z, Dong J, Zhu J-K. 2014. Overproduction of stomatal lineage cells in Arabidopsis mutants defective in active DNA demethylation. Nature Communications 2014 5:1 5, 1–7.

Zhong S, Fei Z, Chen YR, et al. 2013. Single-base resolution methylomes of tomato fruit development reveal epigenome modifications associated with ripening. Nature Biotechnology 31, 154–159.

Zhu JK. 2009. Active DNA demethylation mediated by DNA glycosylases. Annual Review of Genetics 43, 143–166.

