## Supplementary Figure 1, 2 and 3 for "CRISPR/Cas9 gene editing uncovers the role of CTR1 and ROS1 in melon fruit ripening and epigenetic regulation"

**Supplementary Figure 1:** Editions obtain for *ROS1* and *CTR1* in T0 melon plats.

**Supplementary Figure 2:** Phylogenetic tree of the homologous proteins ROS1 in Arabidopsis, tomato and melon.

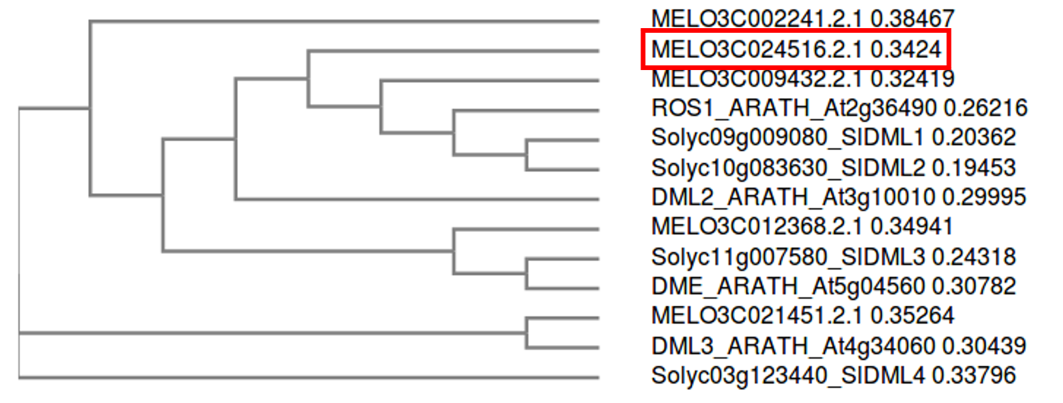

**Supplementary Figure 3:** RNASeq dataset of ROS1 orthologue genes (MELO3C021451, MELO3C024516, MELO3C009432, MELO3C002241) in the climacteric genotype VED and a non-climacteric genotype ¨Piel de sapo (PS)¨.

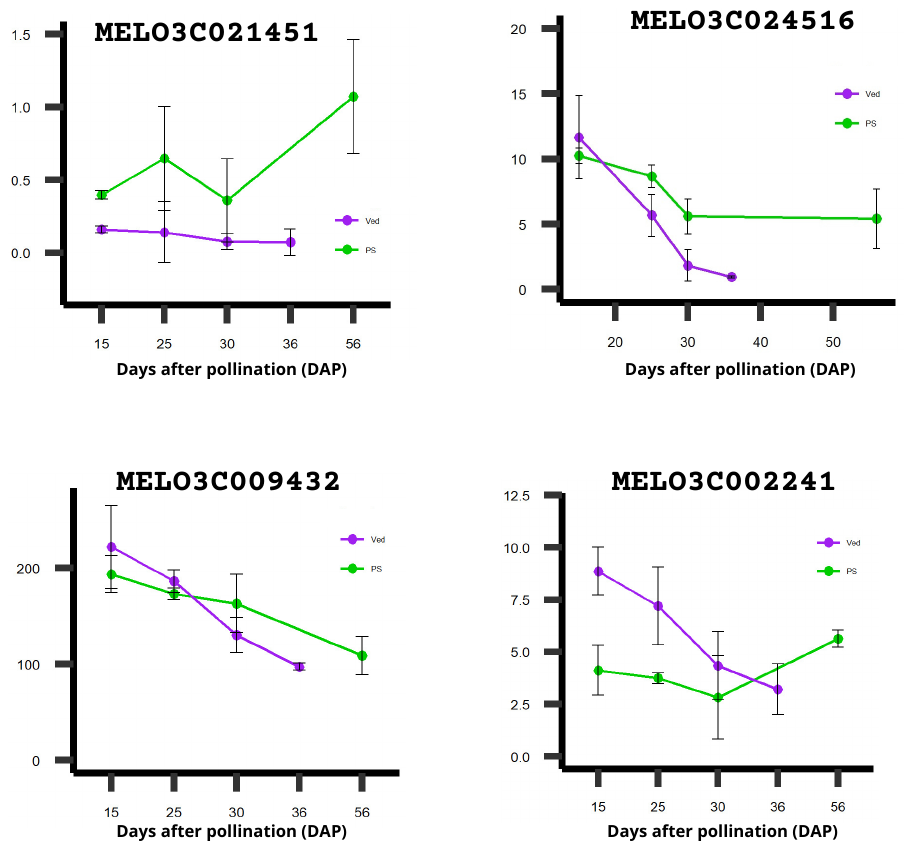
