## Supplementary Table 1, 2 and 3 for "CRISPR/Cas9 gene editing uncovers the role of CTR1 and ROS1 in melon fruit ripening and epigenetic regulation"

**Supplementary information**

**Sup Table 1: List of gRNAs to target *CmROS1* and *CmCTR1-like* genes**

| **Gene target** | **sequence 5´-3´** |
| --- | --- |
| *CmROS1* | gRNA1: AACATCTAGTCCAGCTAGTC |
|  | gRNA2: AACAGGCGCATCGACTGATA |
| *CmCTR1-like* | gRNA: GGGGTCGACTGTACACTATA |

**Sup Table 2: List of primers to detect Cas9 and mutations in CRISPR lines**

| **Gene** | **Primer sequence 5´-3´** |
| --- | --- |
| *CmROS1* | F:CCTCCAGCAGGATGTTTTTG |
|  | R:CACTGCGTCATTTTCATTGC |
| *CmCTR1-like* | F:GCATTCTCCGTCTATCAAAGG |
|  | R:GCTCCGAATCTTCAGAACCA |
| *Cas9* | F:GGACACTTCCTCATCGAGGGT |
|  | R:GTGGAGCCTTGGTGATCTCGG |

**Sup Table 3: Climacteric ripening related traits in two consecutive summer seasons.**

| **Ripening traits (2020)** | VED | | ROS1-CRISPR-1 | | ROS1-CRISPR-2 | | CTR1-CRISPR-1 | |
| --- | --- | --- | --- | --- | --- | --- | --- | --- |
|  | Mean | SD | Mean | SD | Mean | SD | Mean | SD |
| Earliness of production of aroma (EARO) (DAP) | 32,3 | 1,4 | 30,2* | 1 | 30,4* | 1,8 | 30,0* | 1,4 |
| Earliness of chlorophyll degradation (ECD) (DAP) | 32,8 | 1,4 | 30,7* | 1,1 | 30,9* | 1,7 | 30,1* | 1,4 |
| Earliness of abscission layer formation (EALF) (DAP) | 32,8 | 1,4 | 30,5* | 1,2 | 30,7* | 1,9 | 30,1* | 1,4 |
| Flesh firmness (kg · cm^-2^) | 1,6 | 0,9 | 1,9 | 1,2 | 2,1 | 0,9 | 1,6 | 0,5 |
| Maximum ethylene production (µL eth·kg^-1^·h^-1^) | 43,9 | 15,9 | 132,1 | 116 | 119,2 | 55,5 | 55 | 32,2 |
| Earliness of ethylene production (DAPE) (DAP) | 30,2 | 1,3 | 28,8 | 0,8 | 28 | 1,4 | 26,8* | 1,4 |
| Earliness of ethylene peak (DAPP) (DAP) | 31,4 | 1,3 | 30 | 0,7 | 30 | 1,4 | 28,6* | 1,1 |
| Width of ethylene peak (WEP) (days) | 1,8 | 0,4 | 1,2 | 0,4 | 1,4 | 0,5 | 1,2 | 0,4 |

*pvalue <0,05

| **Ripening traits (2021)** | VED | | ROS1-CRISPR-1 | | ROS1-CRISPR-2 | | CTR1-CRISPR-1 | | CTR1-CRISPR-2 | |
| --- | --- | --- | --- | --- | --- | --- | --- | --- | --- | --- |
|  | Mean | SD | Mean | SD | Mean | SD | Mean | SD | Mean | SD |
| Earliness of production of aroma (EARO) (DAP) | 38,0 | 1,3 | 34,3* | 1,4 | 35,0* | 1,7 | 34,8* | 0,8 | 34,4* | 1,9 |
| Earliness of chlorophyll degradation (ECD) (DAP) | 38,0 | 1,3 | 34,2* | 1,2 | 35,2* | 1,5 | 35,0* | 0,6 | 34,6* | 1,5 |
| Earliness of abscission layer formation (EALF) (DAP) | 38,2 | 1,2 | 33,8* | 1,2 | 34,5* | 1,2 | 34,5* | 1,0 | 33,9* | 1,5 |
| Flesh firmness (kg · cm^-2^) | 1,4 | 0,6 | 1,8 | 0,8 | 2,4 | 0,9 | 1,1 | 0,5 | 0,9 | 0,2 |

*pvalue <0,05
